# Identification of a novel transcriptome signature for predicting the response to anti-TNF-α treatment in rheumatoid arthritis patients

**DOI:** 10.1101/2025.05.05.652166

**Authors:** Lucia Santiago-Lamelas, Patricia Castro-Santos, Enrique J. deAndrés-Galiana, Juan Luis Fernández-Martínez, Alejandro Escudero-Contreras, Carlos Pérez-Sanchez, Ismael Sánchez-Pareja, Chary López-Pedrera, Scott A Jelinsky, Raquel Dos Santos Sobrín, Antonio Mera, Josefina Durán, Roberto Díaz-Peña

## Abstract

**Objectives:** To identify and validate a transcriptomic signature capable of predicting the response to antitumour necrosis factor (TNF) therapy in patients with rheumatoid arthritis (RA) before treatment initiation.

**Methods:** We performed a retrospective transcriptomic analysis using two public datasets, RNA-seq data from peripheral blood mononuclear cells in GSE138746 and microarray data from whole blood in GSE33377, to define a small-scale gene signature predictive of the response to anti-TNF treatment. Two external validations were then conducted using data from the COMBINE and the Reina Sofia Hospital cohorts, resulting in a total of 253 individuals, 155 responders and 98 nonresponders.

**Results:** Initial RNA-seq analysis (GSE138746) revealed 53 genes that were differentially expressed between responders and nonresponders; however, none of these genes remained significant after p value adjustment with the Benjamini–Hochberg method. A small-scale genetic signature comprising the 18 most discriminatory genes was then developed, achieving a leave-one-out cross-validation (LOOCV) predictive accuracy of 88.75%. We further refined this list to seven genes (*COMTD1, MRPL24, DNTTIP1, GLS2, GTPBP2, IL18R1*, and *KCNK17*) that effectively predicted the response to anti-TNF-α treatment, with an area under the receiver operating characteristic (ROC) curve (AUC) of 0.84 in the GSE33377 dataset. Internal validation of the GSE138746 dataset yielded an AUC of 0.89. Finally, external validation confirmed the robustness of the seven-gene model (AUC≥ 0.85).

**Conclusions:** We identified a transcriptomic signature that aids the prediction of the response to anti-TNF treatment in RA patients. These findings support its potential use as a precision medicine tool to improve therapeutic decision-making and reduce exposure to ineffective treatments in RA patients.

## INTRODUCTION

Rheumatoid arthritis (RA) is a chronic, systemic autoimmune disease affecting approximately 0.5–1% of the global population, with a higher prevalence in females than in males. It is characterised by inflammation of the synovial joints, leading to cartilage and bone destruction, functional disability, and a significant reduction in patients’ quality of life (1). Sequelae are frequent, and up to half of patients are unable to work within ten years after diagnosis (2,3).

Despite advances in treatment, the management of RA remains a challenge because of its heterogeneity and variable responses to available therapies. Currently, all RA patients must receive disease-modifying antirheumatic drugs (DMARDs) as part of their pharmacologic treatment (4). These drugs can be classified into conventional, targeted, and biological DMARDs. Among the biological DMARDs, tumour necrosis factor (TNF)-α inhibitors were the first to be developed. Anti-TNF agents have revolutionised the treatment of RA, significantly improving clinical outcomes and inhibiting radiographic progression in a substantial proportion of patients (5). However, although anti-TNF therapy is effective in most cases, up to 30% of treated patients show an inadequate response (6). The mechanisms behind this variability are multifactorial, encompassing genetic, epigenetic, and environmental factors, and remain poorly understood (7). This high heterogeneity in the response to therapy underscores the need to reveal the pathophysiological mechanisms of the disease to optimise treatment outcomes. Moreover, the high cost of biological therapies, particularly in chronic conditions such as RA, poses a significant challenge to the sustainability of health care systems worldwide. Given the economic impact and potential for adverse effects, enhancing treatment efficiency by optimising the selection of patients who are most likely to benefit from anti-TNF therapy is imperative.

Robust molecular biomarkers are needed in clinical practice to guide anti-TNF therapy in RA patients. Numerous transcriptomic studies aimed at deciphering the biological processes associated with anti-TNF responses have been conducted, analysing both the synovium and blood of RA patients (8–16). Although these studies demonstrated the potential of transcriptome profiling to predict the response to anti-TNF therapy before treatment initiation, a consistent reproduction of results across different studies has not yet been achieved.

In the present study, we hypothesised that we would increase the predictive power for treatment response in patients with RA by integrating multiple gene expression profiles through bioinformatic analyses. Using this approach, we identified a transcriptomic signature capable of accurately predicting the response to anti-TNF therapy prior to treatment initiation.

## MATERIALS AND METHODS

### Data sources and patients

The study scheme for uncovering and testing the capacity of a transcriptomic signature to predict the anti-TNF response is shown in Figure 1. We performed a retrospective analysis of whole peripheral blood mononuclear cell (PBMC) transcriptomic profiles from patients with RA to define the small-scale signature genes involved in the response to anti-TNF-α treatment. We defined a small-scale signature as the minimal subset of genes capable of maximising cross-validation accuracy when predicting treatment response (responder vs. nonresponder). This parsimony-based approach assumes that a compact gene set can best capture the relevant low-frequency features in high-dimensional gene expression data, minimising model overfitting and accounting for inherent uncertainty in biological systems. We accessed the publicly available dataset GSE138746 from the Gene Expression Omnibus (GEO) database (https://www.ncbi.nlm.nih.gov/geo/) (17). This dataset contained 80 samples with gene expression profiles of PBMCs, including 38 and 42 samples from RA patients treated with adalimumab (ADA) and etanercept (ETN), respectively. An additional publicly available dataset (GSE33377) profiled by microarray was downloaded from the GEO database to define, based on the small-scale signature genes, a transcriptomic signature capable of predicting response prior to anti-TNF treatment. The GSE33377 dataset contained expression profiles of the whole blood of 42 RA patients starting treatment with infliximab (IFX) or ADA (13).

**Figure 1.**
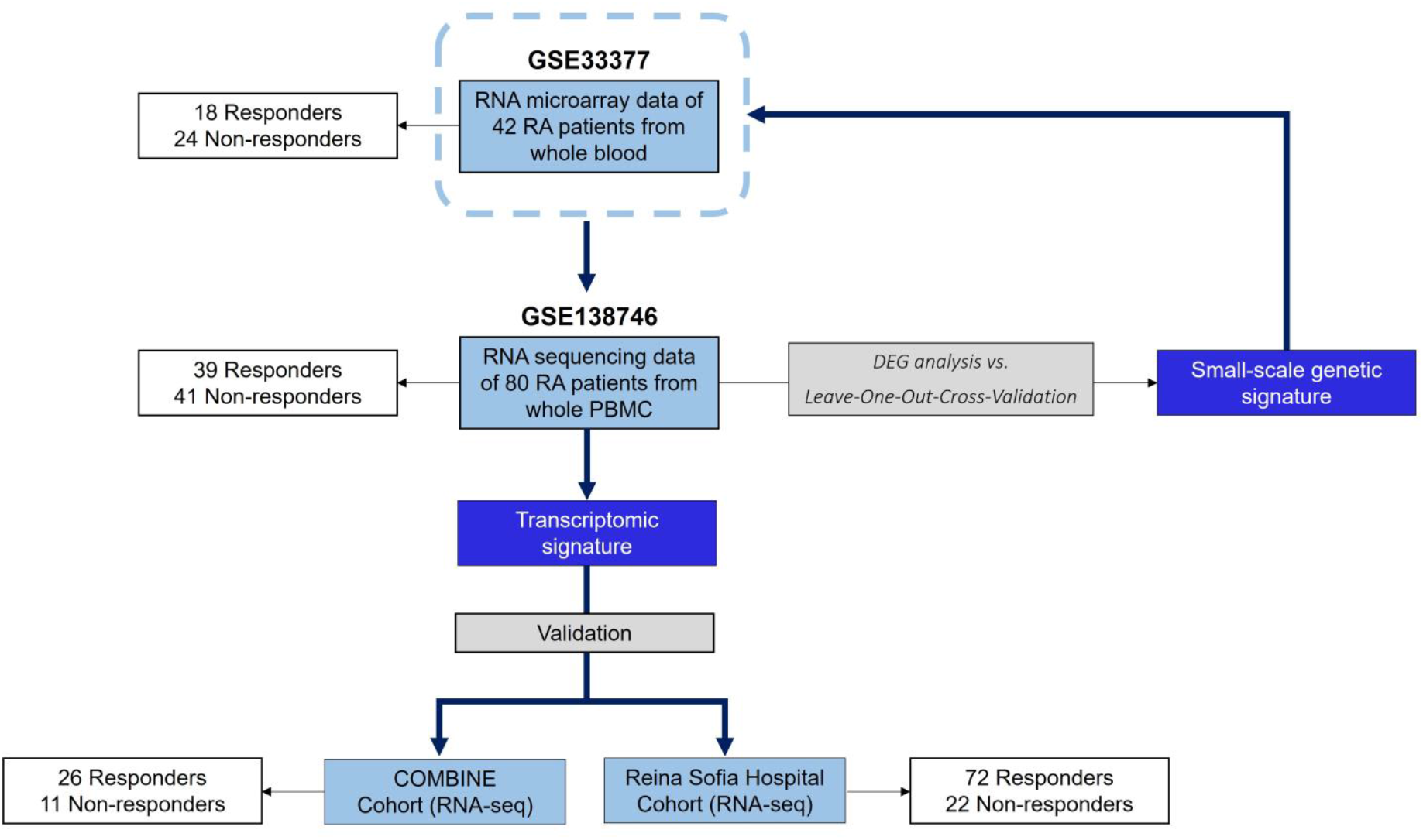
Flowchart utilised to search for a transcriptomic signature that predicts the response to anti-TNF treatment. Abbreviations: DEGs, differentially expressed genes; PBMCs, peripheral blood mononuclear cells; RA, rheumatoid arthritis.

For independent validation of the predictive signature, we used two independent sets of samples. The Reina Sofia Hospital cohort included 94 RA patients with moderate to high disease activity who were recruited at the Reina Sofia Hospital from Cordoba, Spain. These patients started TNF therapy after failing standard DMARD treatment, and the cohort included 72 responders and 22 nonresponders (Table 1). Additionally, we included the COMBINE cohort (18), comprising 37 individuals who previously did not respond to methotrexate (MTX) treatment and who initiated TNF inhibitors (TNFi) therapy. This cohort included 26 TNFi responders (R) and 11 nonresponders (NR). In both the COMBINE and the Reina Sofia Hospital cohorts, good and moderate EULAR responders were classified as “responders” and were compared with the EULAR “nonresponders.”

**Table 1.** Clinical characteristics of the Reina Sofía Hospital RA cohort.

| Clinical parameters | RA patients (n=94) |
| --- | --- |
| Gender (Female/Male) | 73/21 |
| Age, years (mean ± SD) | 53.91 ± 10.99 |
| Disease evolution, years (mean ± SD) | 9.38 ± 8.38 |
| TJC (mean ± SD) | 6.24 ± 5.29 |
| SJC (mean ± SD) | 3.81 ± 4.07 |
| DAS28-CRP (mean ± SD) | 4.43 ± 1.09 |
| DAS28-ESR (mean ± SD) | 4.51 ± 1.31 |
| SDAI (mean ± SD) | 24.80 ± 11.08 |
| CDAI (mean ± SD) | 23.36 ± 10.81 |
| Smoking (n, %) | 29/93 (31.18%) |
| Arterial hypertension (n, %) | 25/91 (27.47%) |
| Diabetes (n, %) | 5/93 (5.38%) |
| Extra-articular manifestations (n, %) | 30/91 (32.97%) |
| Radiological involvement (n, %) | 52/79 (65.82%) |
| Eroded joints (mean ± SD) | 1.39 ± 2.41 |
| **Laboratory parameters** |  |
| CRP, mg/mL (mean ± SD) | 14.43 ± 19.78 |
| ESR, mm/h (mean ± SD) | 21.36 ± 19.27 |
| Anti-CCPs, IU/mL (mean ± SD) | 159.92 ± 199.46 |
| RF, IU/mL (mean ± SD) | 175.88 ± 257.89 |
| **Treatments** |  |
| NSAIDs (n, %) | 75/94 (79.79%) |
| Corticosteroids (n, %) | 81/94 (86.17%) |
| Methotrexate (n, %) | 56/94 (59.57%) |
| Leflunomide (n, %) | 32/94 (34.04%) |
| Hydroxychloroquine (n, %) | 28/94 (29.79%) |
| Sulfasalazine (n, %) | 5/94 (5.32%) |

Abbreviations: Anti-CCP, anti-cyclic citrullinated peptide; CDAI, Clinical Disease Activity Index; CRP, C-reactive protein; DAS28-CRP, Disease Activity Score 28 based on C-reactive protein; DAS28-ESR, Disease Activity Score 28 based on erythrocyte sedimentation rate; ESR, Erythrocyte sedimentation rate; NSAIDs, Non-steroidal anti-inflammatory drugs; RA, Rheumatoid arthritis; RF, Rheumatoid factor; SDAI, Simplified Disease Activity Index; SJC, Swollen joint count; TJC, Tender joint count; Vit D, Vitamin D.

### Analysis of differentially expressed genes

To perform an initial analysis of differential expression, the statistical analyses were performed using R version 4.1.1. After the expression matrix of the GSE138746 dataset was processed via the DESeqDataSetFromMatrix R function, principal component analysis and differential expression analysis were performed with the DESeq function (19). Differential gene expression analysis was performed via the parametric Wald test with the Benjamini—Hochberg adjustment method. Genes with counts with means lower than 5 were removed from the DESeq table before p value adjustment. The GSEA algorithm was used for enrichment analysis of Wikipathways and GO terms (cellular component, biological processes and molecular function) (20).

### Genetic small-scale signature

The identification of a transcriptomic signature capable of predicting response prior to anti-TNF treatment constitutes a phenotype prediction problem. This category of problems is characterised by a high degree of underdeterminacy, given that the number of monitored genes far exceeds the number of samples employed. It is imperative to prioritise genes based on their discriminatory power in predicting the phenotype and to focus on sampling highly predictive networks anticipated to be involved in genetic pathways underlying treatment response.

These networks occupy the uncertainty space linked to the classifier used for phenotype discrimination, a structure previously discussed in the context of linear and nonlinear inverse problems (21,22). The smallest-scale signature within a classifier includes the fewest genes with optimal predictive accuracy. However, noise in genetic data means that some high-accuracy signatures may not align with relevant genetic pathways (23,24). Addressing parameter identification thus requires accounting for noise, highlighting the need to evaluate the discriminatory power of key phenotype-related genes using sampling methods.

In this study, we applied Fisher’s ratio sampler (25) to identify defective pathways on the basis of genes with high discriminatory power, as indicated by Fisher’s ratio. Genes with the highest Fisher’s ratios among those with significant fold changes were selected, with a focus on differentially expressed genes in both the over- and underexpressed tails. A cut-off of 0.8 for the Fisher ratio defines the set of discriminatory genes, which is adjustable to 0.5 if needed when the number of such genes is limited. To establish a small-scale genetic signature, genes are ranked by discriminatory power, and recursive feature elimination identifies the smallest optimal signature. The predictive accuracy is assessed via leave-one-out cross-validation (LOOCV) with a nearest neighbour classifier, which approximates the typical size of high-discriminatory networks.

The random sampler identifies alternative networks of highly discriminatory genes, assigning each gene a sampling probability proportional to its Fisher’s ratio. Networks are randomly constructed on the basis of this distribution, and their LOOCV predictive accuracy is evaluated. This process follows Bayes’ rule, incorporating a prior probability based on the Fisher’s ratio of selected genes and a likelihood function dependent on the network’s LOOCV accuracy.

Ultimately, considering the most discriminatory networks with a predictive accuracy exceeding a predefined threshold (typically greater than 85%), subsequent sampling frequencies of the primary prognostic genes involved in these networks are determined.

### Predictive model for TNFi treatment response

To construct a predictive model for TNFi treatment response, logistic regression was employed, incorporating the normalised expression levels of seven genes (*KCNK17, DNTTIP1, IL18R1, GLS2, MRPL24, GTPBP2*, and *COMTD1*), interaction terms, and demographic variables (sex and age) as predictors. The predicted probabilities of response were then calculated for each patient, and the model’s discriminative performance was evaluated using ROC curve analysis and the AUC.

### Statistical analysis

Statistical analyses were performed using R version 4.2.1. The following R packages were used: tidyverse (26), pROC (27), ggplot2 (28), dplyr (29), caret (30), leaps (31), patchwork (32), and ggpubr (33). For boxplot comparisons of gene expression between R and NR, p values were calculated using the Wilcoxon rank-sum test. ROC curves were generated using the pROC package, and AUC values were compared using DeLong’s test for statistical significance.

## RESULTS

### Analysis of differentially expressed genes in the GSE138746 dataset

An initial analysis of the RNA-seq results revealed 53 genes that were differentially expressed between R and NR, including 28 upregulated and 25 downregulated genes in R versus NR, according to the following criteria: fold change > |1| and *p* value <0.05. To visualise the differentially expressed genes, we constructed a volcano plot (Figure 2A). No significant differences were found between the groups when the p value was adjusted by the Benjamini–Hochberg adjustment method (Supplementary Table 1). Moreover, the representation of the heatmap and the principal component analysis (PCA) analysis revealed poor separation of the samples from R and NR to anti-TNF therapy according to the 53 differentially expressed genes identified via RNA-seq analysis (Figure 2B and 2C, respectively). The functional enrichment analysis did not reveal any significant relationship between the analysed genes and the response (all the adjusted p values>0.1; Supplementary Tables 2-5).

**Figure 2.**
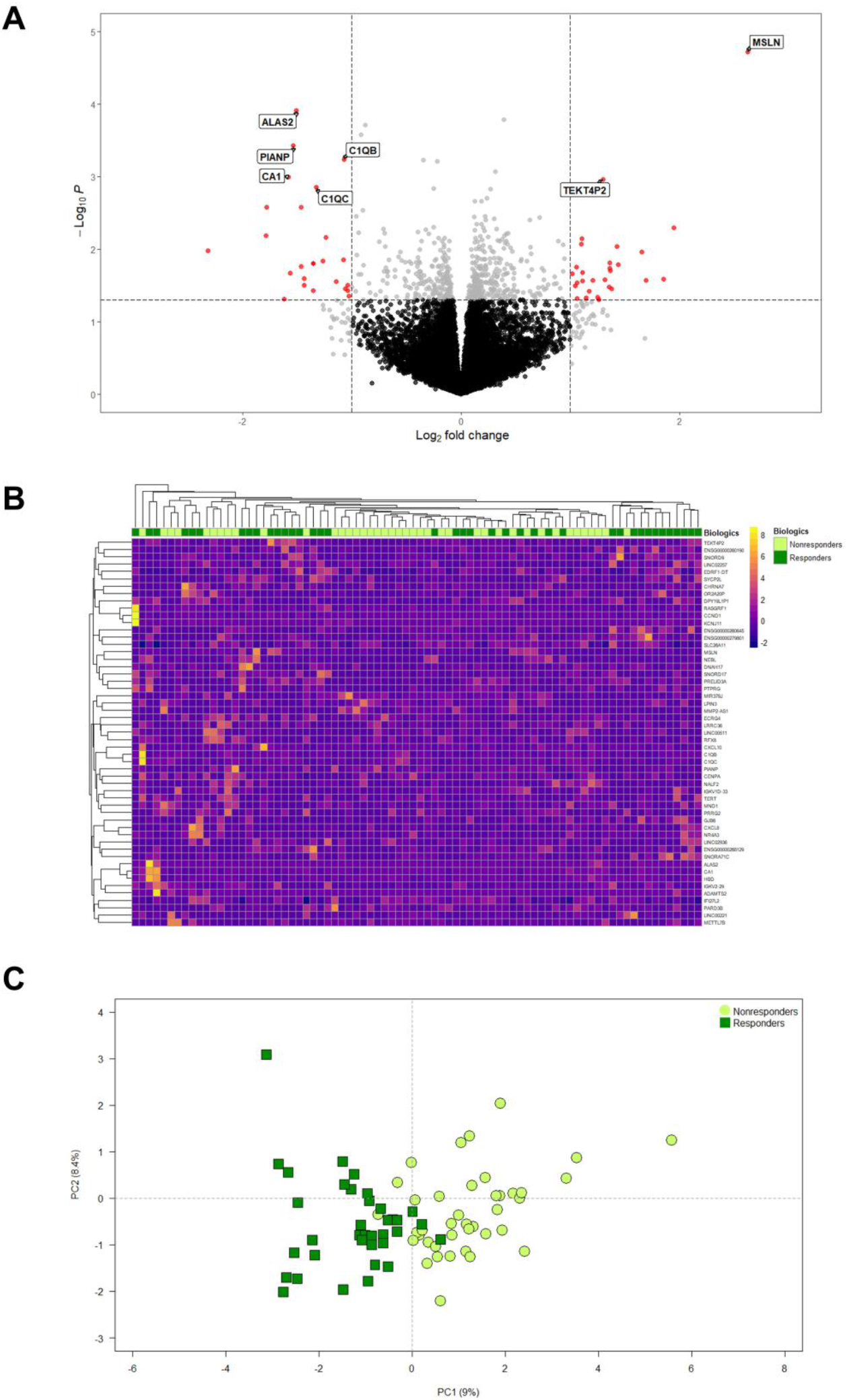
Differential expression analysis of the GSE138746 dataset. (**A**) Volcano plot of differentially expressed genes (fold change > |1| and *p* value <0.05); (**B**) heatmap and (**C**) principal component analysis of 53 differentially expressed genes at baseline (fold change > |1| and *p* value <0.05).

### Identification of a highly discriminatory gene set

Table 2 shows the list of the most discriminatory genes (calculated with Fisher’s ratio sampler) associated with the phenotypes of the R patients compared with the NR patients obtained through Fisher’s ratio sampler once. The small-scale genetic signature identified was composed of the 18 most discriminatory genes (*KCNK17, LOC100506235, TRBV6-4, GLS2, MRPL24, DNTTIP1, MYPOP, LPIN3, OR2A9P, GTPBP2, ZFTA, COMTD1, ZFP57, IL18R1, KRBOX5, FEM1C, ENSG00000278627*, and *THBS4-AS1*), with a LOOCV predictive accuracy of 88.75%.

**Table 2.** Most discriminatory genes were identified via Fisher’s ratio sampler.

| Gene | MedianC1 | StdC1 | MedianC2 | StdC2 | FC | FR | LOOCV |
| --- | --- | --- | --- | --- | --- | --- | --- |
| KCNK17 | 1.0 | 13.28 | 15.0 | 13.04 | -0.92 | 0.57 | 67.50 |
| LOC100506235 | 29.0 | 15.90 | 13.0 | 14.49 | 0.64 | 0.55 | 68.75 |
| TRBV6-4 | 12.0 | 12.45 | 0.0 | 11.71 | 0.93 | 0.49 | 70.00 |
| GLS2 | 68.0 | 37.01 | 35.0 | 30.20 | 0.53 | 0.48 | 68.75 |
| MRPL24 | 456.0 | 107.41 | 356.0 | 110.06 | 0.20 | 0.42 | 72.50 |
| DNTTIP1 | 639.0 | 197.64 | 464.0 | 183.20 | 0.27 | 0.42 | 72.50 |
| MYPOP | 101.0 | 51.91 | 150.0 | 56.29 | -0.33 | 0.41 | 78.75 |
| LPIN3 | 12.0 | 16.64 | 0.0 | 9.49 | 1.22 | 0.39 | 73.75 |
| OR2A9P | 16.0 | 12.90 | 4.0 | 15.38 | 0.20 | 0.36 | 75.00 |
| GTPBP2 | 1824.0 | 341.08 | 1539.0 | 337.47 | 0.12 | 0.35 | 70.00 |
| ZFTA | 141.0 | 56.53 | 189.0 | 58.65 | -0.20 | 0.35 | 76.25 |
| COMTD1 | 199.0 | 65.24 | 150.0 | 51.84 | 0.37 | 0.35 | 77.50 |
| ZFP57 | 7.0 | 67.25 | 72.0 | 89.25 | -0.59 | 0.34 | 81.25 |
| IL18R1 | 377.0 | 136.53 | 508.0 | 181.10 | -0.36 | 0.33 | 86.25 |
| KRBOX5 | 324.0 | 81.78 | 398.0 | 99.33 | -0.20 | 0.33 | 86.25 |
| FEM1C | 597.0 | 166.85 | 764.0 | 242.94 | -0.20 | 0.32 | 86.25 |
| ENSG00000278627 | 4.0 | 9.79 | 13.0 | 12.51 | -0.78 | 0.32 | 86.25 |
| THBS4-AS1 | 48.0 | 27.61 | 27.0 | 25.11 | 0.52 | 0.32 | 88.75 |
| ECRG4 | 16.0 | 15.97 | 5.0 | 11.49 | 1.02 | 0.31 | 86.25 |
| ENSG00000215447 | 120.0 | 48.23 | 84.0 | 43.71 | 0.32 | 0.31 | 82.50 |
| LRRC75A | 98.0 | 39.62 | 68.0 | 37.10 | 0.28 | 0.31 | 78.75 |
| NDUFA8 | 340.0 | 102.98 | 262.0 | 97.07 | 0.22 | 0.30 | 83.75 |
| RPS24P19 | 15.0 | 6.14 | 10.0 | 6.68 | 0.31 | 0.30 | 82.50 |
| COPG2 | 526.0 | 150.26 | 419.0 | 124.08 | 0.29 | 0.30 | 81.25 |

The median C1 and median C2 represent the median values of each group (C1 represents the response to anti-TNF-α treatment, and C2 represents the nonresponse). Abbreviations: FC, fold change value; FR, fisher's ratio value; LOOCV, mean accuracy in Leave-One-Out-Cross-Validation; Std, standard deviation of each group.

### A transcriptomic signature was associated with the prediction of response to anti-TNF therapy

Next, we evaluated the discriminatory power of the 18 most discriminatory genes to create a transcriptomic signature capable of accurately distinguishing R from NR using ROC analyses. Using a logistic regression model, we selected a predictive model of 7 genes (*MRPL24, COMTD1, DNTTIP1, GLS2, GTPBP2, IL18R1*, and *KCNK17*) that effectively discriminated R and NR (AUC=0.84; Figure 3). Single genes showed a low to moderate discriminatory power to distinguish between R and NR (AUC= 0.43-0.78, Supplementary Figure 1).

**Figure 3.**
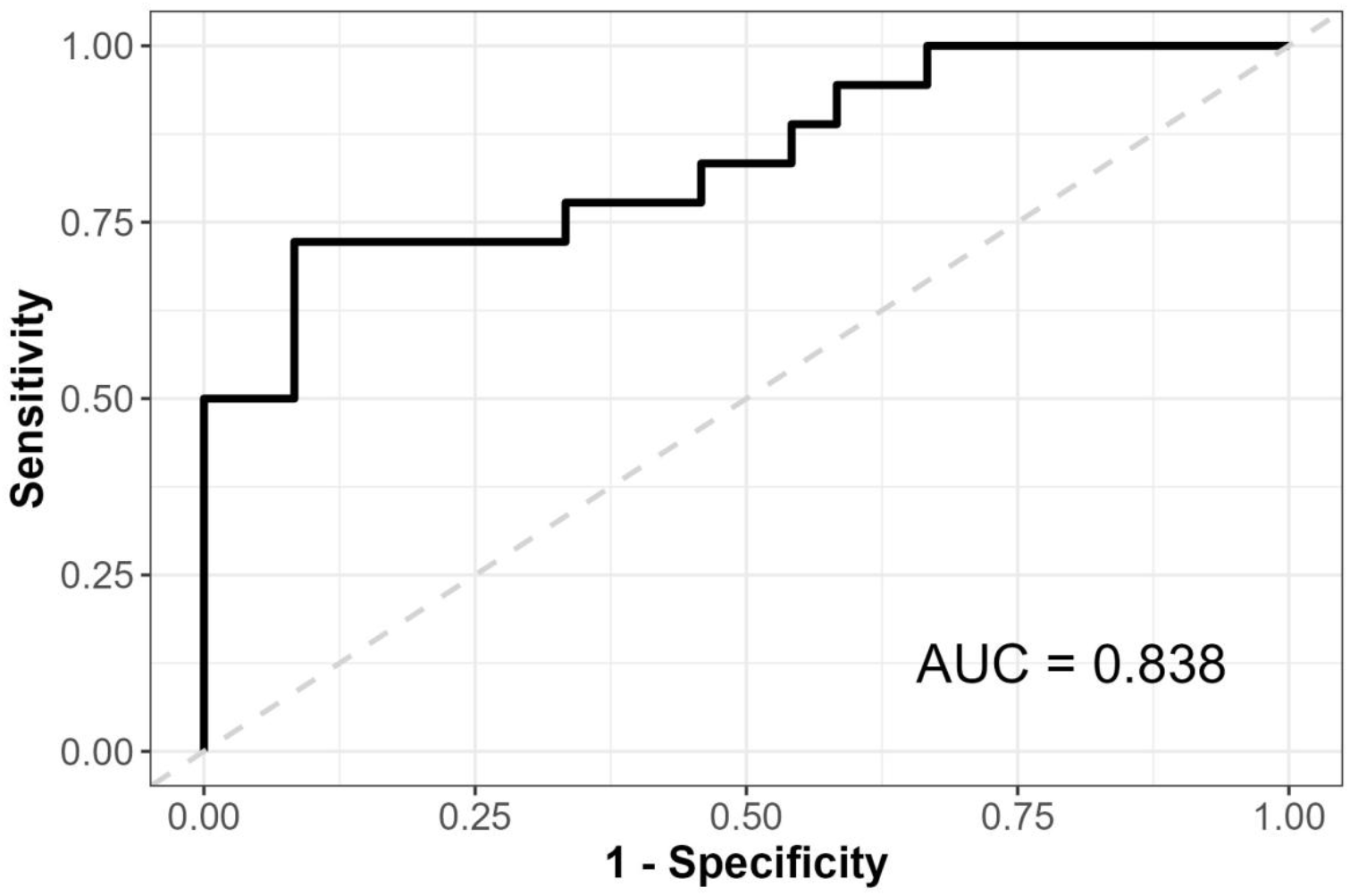
ROC curve for predicting response using the transcriptomic signature in the GSE33377 dataset.

We then performed a validation using the first dataset (GSE138746), the same dataset we used to establish the genetic small-sale signature. Boxplots for each gene revealed significant differences in expression levels between R and NR (p value<0.05 for *GLS2, MRPL24, GTPBP2*, and *DNTTIP1* and p value<0.005 for *KCNK17, IL18R1*, and *COMTD1*). ROC analysis of the 7 genes revealed high predictive performance, with an AUC of 0.949 (Figure 4A). Additionally, we evaluated the predictive performance of the transcriptomic signature in comparison with that of conventional serological biomarkers. The 7-gene model significantly outperformed both anti-cyclic citrullinated peptide (anti-CCP) and rheumatoid factor (RF) in terms of discriminative capacity, achieving AUCs of 0.949 versus 0.519 and 0.555, respectively (Table 3).

**Table 3.** Comparative performance metrics of the transcriptomic signature versus anticyclic citrullinated peptide and rheumatoid factor in the GSE138746 dataset.

| **Biomarker** | **AUC** | ***p-value*** | **Sensitivity** | **Specificity** | **PPV** | **NPV** |
| --- | --- | --- | --- | --- | --- | --- |
| Transcriptomic Signature | 0.949 | $6.14 \times 10^{-75}$ | 0.872 | 0.951 | 0.944 | 0.886 |
| Anti-CCP | 0.519 | 0.701 | 0.769 | 0.268 | 0.500 | 0.550 |
| RF | 0.555 | 0.293 | 0.744 | 0.366 | 0.527 | 0.600 |

Abbreviations: Anti-CCP, anti-cyclic citrullinated peptide; AUC, Area under the curve; NPV, Negative predictive value; PPV, Positive predictive value; RF, rheumatoid factor.

**Figure 4.**
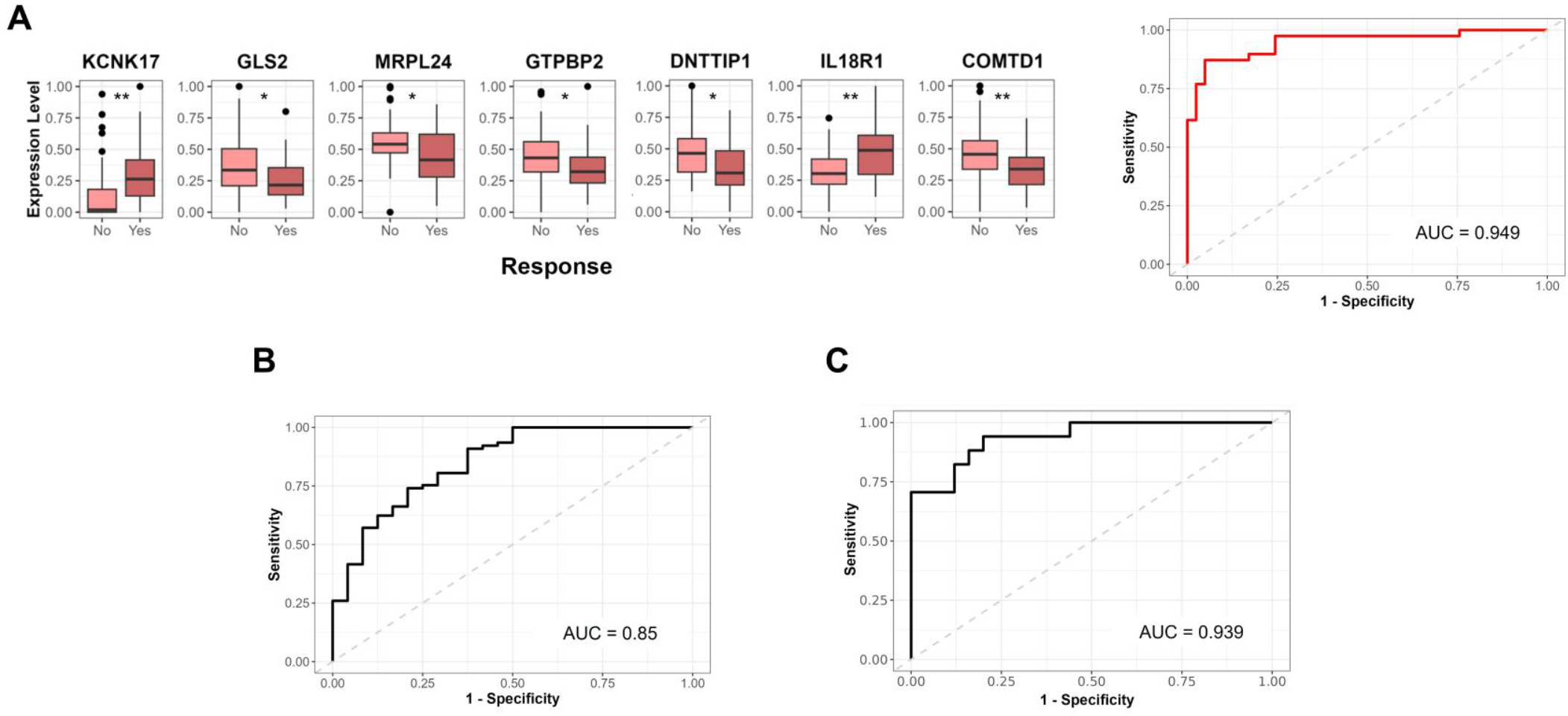
Validation of the 7-gene transcriptomic signature for predicting the response to anti-TNF therapy. (**A**) Boxplots showing the expression levels of each gene in the 7-gene signature (KCNK17, GLS2, MRPL24, GTPBP2, DNTTIP1, IL18R1, and COMTD1) between responders (R) and nonresponders (NR) in the GSE138746 dataset. Statistical significance was assessed using the Wilcoxon rank-sum test (\**p value* < 0.05; \*\**p value* < 0.005). The ROC curve displays the discriminatory power of the 7-gene signature in the GSE138746 dataset; (**B**) ROC curve for the 7-gene transcriptomic signature applied to the Reina Sofía Hospital cohort and (**C**) ROC curve for the COMBINE cohort.

To validate the signature, we applied the model to two independent datasets. The first validation cohort achieved an AUC of 0.85, supporting the robustness of the 7-gene model in distinguishing R from NR to anti-TNF therapy (The Reina Sofía Hospital Cohort, Figure 4B). The second dataset, the COMBINE cohort (18), confirmed the predictive value of the signature, with an AUC of 0.939 (Figure 4C).

## DISCUSSION

One of the major challenges in the treatment of RA is understanding the biological mechanisms that influence the clinical response to anti-TNF therapy. Despite the transformative impact of TNF inhibitors on RA treatment, a significant proportion of patients exhibit partial response or nonresponse, leading to considerable clinical and economic burdens. The complexity of the pathophysiology of RA, combined with patient-specific genetic, environmental, and lifestyle factors, contributes to the observed variability in treatment outcomes. This heterogeneity highlights the urgent need for predictive biomarkers to guide individualised treatment strategies and ensure that patients receive the most effective interventions early in the disease course. Historically, research on predictive markers of the anti-TNF response has been limited by the range of techniques and analytical methods used, leading to single-gene analyses or potential biomarker analyses without considering the broader molecular landscape. The advent of high-throughput sequencing technologies and the application of advanced bioinformatics algorithms have provided an unprecedented opportunity to dissect the molecular complexity underlying therapeutic resistance.

Our findings suggest that the transcriptomic signature composed of the genes *COMTD1, MRPL24, DNTTIP1, GLS2, GTPBP2, IL18R1*, and *KCNK17* is a powerful tool for predicting the response to anti-TNF treatment in RA patients. The robustness of this signature is supported by external validation in the COMBINE and Reina Sofia Hospital cohorts, indicating its applicability across different clinical settings.

The *KCNK17* gene is located on chromosome 6 and encodes a member of the potassium two-pore domain channel subfamily K member 17. While the direct association between potassium channels and RA is not well established, some studies suggest that proper dietary potassium intake may play a significant role in reducing pain and inflammation (34). The *GLS2* gene encodes a mitochondrial phosphate-activated glutaminase that is predominantly expressed in the liver and associated with the tumour suppressor protein p53. The inhibition of GLS2 has been shown to enhance the process of ferroptosis—a type of iron-dependent cell death—linked to various autoimmune diseases, including RA, by amplifying the inflammatory response (35). Furthermore, recent studies have indicated that excessive iron intake can lead to the degradation and destruction of the cartilage matrix (36). In addition, GLS2 has been shown to support IL-2 production by CD4+ T cells, suggesting a role in modulating immunity (37). The *GTPBP2* gene encodes GTP binding protein 2, which is involved in protein synthesis elongation and various signal transduction pathways. In fact, GTPases are crucial in many complex cellular processes. In particular, small GTPases play important roles as controllers of neutrophil—endothelial cell interactions and are related to neutrophil recruitment to inflammation points. However, the biological functions in which GTPBP2 is involved have not yet been fully characterised (38). The *DNTTIP1* gene, located on chromosome 20, encodes deoxynucleotidytransferase terminal interacting protein 1 (DNTTIP1). DNTTIP1 is involved in gene transcription and cell proliferation and has been associated with immune infiltration, a typical feature of RA (39). The *IL18R1* gene encodes the receptor of the cytokine IL-18, which is involved in the response to infection, inflammation, and autoimmunity (40). IL-18 enhances immune responses and is also involved in maintaining barrier homeostasis and signalling in inflammatory processes. High levels of IL-18 have been associated with various inflammatory and autoimmune diseases. Although the broad biological roles of IL-18 complicate therapeutic strategies, novel approaches, such as blocking its activation or release, are currently under study (40). The *COMT1* gene encodes an O-methyltransferase enzyme, the biological function of which is still poorly understood. The *COMTD1* gene has been identified as a part of a diagnostic and therapeutic signature related to cuproptosis and ferroptosis in patients with oesophageal squamous cell carcinoma (41). Given the involvement of ferroptosis in increasing inflammatory responses, the role of COMTD1 in RA may be linked to this mechanism. *MRPL24* (mitochondrial ribosomal protein L24) is a gene located on chromosome 1 that is essential for the proper function of the mitochondrial translation process (42). Mutations in *MRP* genes are involved in conditions such as cancer, neurodegeneration, obesity and inflammatory diseases (43). Several genes within our 7-gene transcriptomic signature have been linked to key intracellular pathways involved in the response to anti-TNF therapy. Recent studies suggest that genetic and molecular factors, including differentially methylated positions (DMPs) in immune-related genes (*COMTD1*) (44), inflammatory cytokine signalling (*IL18R1*) (45), nucleosome autoantigen formation (*DNTTIP1*) (46), and RNA splicing mechanisms (*GTPBP2*) (47), may influence treatment effectiveness. These findings support the biological relevance of our transcriptomic signature in predicting the response to anti-TNF therapy. However, further studies will be essential to validate these relationships and elucidate the precise role of these genes within the molecular pathways involved in the treatment response. A deeper understanding of these mechanisms could provide new insights into disease pathophysiology and potentially guide the development of novel therapeutic strategies for RA and other autoimmune diseases.

In a relevant study, Toonen et al. focused on validating eight previously identified gene expression signatures that predict the response to anti-TNF therapy prior to treatment initiation (13). Although they assessed a panel of 20 genes that achieved a sensitivity of 71% and a specificity of 61%, none of these genes overlapped with those in our signature. In another study, the transcriptome of whole blood cells was measured at two time points, pretreatment and three months after the initiation of ADA therapy, but no statistically significant differences were found between R and NR patients at either time point. (16). However, more recently, a study analysing differential expression and gene methylation revealed clear differences between R to ADA and R to ETN (17), allowing the development of two machine learning models for the prediction of the response to either ADA or ETN using differential genes, with high accuracy (85.9% and 79%, respectively).

Several studies have investigated gene expression profiles in samples from the RA synovium. A 2006 study examined gene expression profiles in arthroscopic biopsies of RA patients before and after treatment with IFX (9). This study identified a distinct set of 279 genes that were differentially expressed between R and NR, highlighting the potential role of the *MMP-3* gene as a predictive marker for favourable treatment response. Similarly, in a study analysing gene expression in the synovium of RA patients treated with ADA, Badot et al. reported that responders presented increased expression of genes associated with pathways critical for cell division and immune response regulation, particularly cytokines, chemokines, and their receptors. In this case, the genes linked to the response to TNFi were the *IL-7R, IL-6, INDO, CDC2, MK167* and *GTSE1* genes, none of which matched the genes described in our signature (11). Additionally, in RNA samples from patient synovium, Aterido et al. revealed differential expression profiles in response to anti-TNF therapy (15), with clear differences in the profiles depending on the type of anti-TNF used.

To our knowledge, this is the first study demonstrating the involvement of this gene signature in predicting the response to anti-TNF therapy. Unlike previous studies that identified genetic signatures predictive of responses to a single type of TNF inhibitor, our work represents a significant advancement by revealing a transcriptomic signature that is consistent across different anti-TNF drugs. This broad applicability underscores the potential of our signature in guiding treatment decisions for a wider patient population, thereby enhancing the personalisation of therapy in RA. The comprehensive search for molecular predictors of anti-TNF responses using advanced computational methods is clearly a field of high interest in RA, representing a promising tool for the decision tree of more adequate therapies to be used. Our work represents a major advance in the field and demonstrates the ability to integrate molecular data with machine learning models to develop precision medicine strategies for RA.

On the other hand, our study had several limitations that preclude solid conclusions. First, its retrospective design may introduce biases in data accuracy, as it relies on preexisting transcriptomic datasets and clinical records. Additionally, the lack of longitudinal transcriptomic data prevents an in-depth evaluation of dynamic gene expression changes throughout the course of treatment, which could further enhance the predictive capacity of the identified signature. Future prospective studies incorporating serial transcriptomic analyses could provide a more comprehensive understanding of treatment response trajectories. Another limitation is the limited ethnic diversity within the studied cohorts, which may restrict the generalisability of our findings to broader populations. Genetic variations across different ethnic groups could influence transcriptomic profiles and treatment responses, highlighting the need for more diverse patient cohorts in future validation studies. Furthermore, while the predictive model was validated across multiple independent datasets and anti-TNF agents, additional validation in larger, real-world clinical settings is necessary to confirm its robustness and clinical applicability. Finally, while our analysis identified a strong transcriptomic signature, it remains essential to integrate other molecular and clinical factors to improve prediction accuracy. The inclusion of multiomics approaches, such as proteomics or metabolomics, along with key clinical and serological biomarkers, could provide a more holistic framework for optimising treatment decisions in RA.

In conclusion, this study provides a useful tool to optimise RA treatment strategies, improving the criteria for treatment selection prior to the start of RA treatment. The gene signature we describe will help to predict the response to anti-TNF agents in RA patients before the initiation of therapy, reducing the time that a patient may spend on ineffective treatment, as well as preventing the potential development of negative side effects.

## ACKNOWLEDGMENTS

Roberto Díaz-Peña is supported by the Miguel Servet (CP21/00003) contract, funded by the Instituto de Salud Carlos III (ISCIII) and co-funded by the European Union. Chary López-Pedrera was supported by a contract from the Spanish Junta de Andalucía (‘Nicolas Monardes’ programme). Carlos Pérez-Sanchez was financed by a contract from MINECO ‘Ramon y Cajal’ program (RYC2021-033828-I), cofounded by the European Union ‘NextGenerationEU’/PRTR.

This work was supported by grants and support from the Fondecyt grant Nº 1220540, ISCIII) and co-funded by the European Union (PI22/00804), and GAIN Proyectos de Excelencia (IN607D2022/06). It was also supported by grants: PI24/00959 funded by ISCIII and co-funded by the European Union; ‘RD24/0007/0019’ _-Health Outcomes-Oriented Cooperative Research Networks-, granted by the “Instituto de Salud Carlos III (ISCIII)” _and “Funded by the European Union – NextGenerationEU”, via “Mecanismo de Recuperación y Resiliencia (MRR)” and “Plan de Recuperación, Transformación y Resiliencia (PRTR)”.

---

**Supplementary Figure 1.**
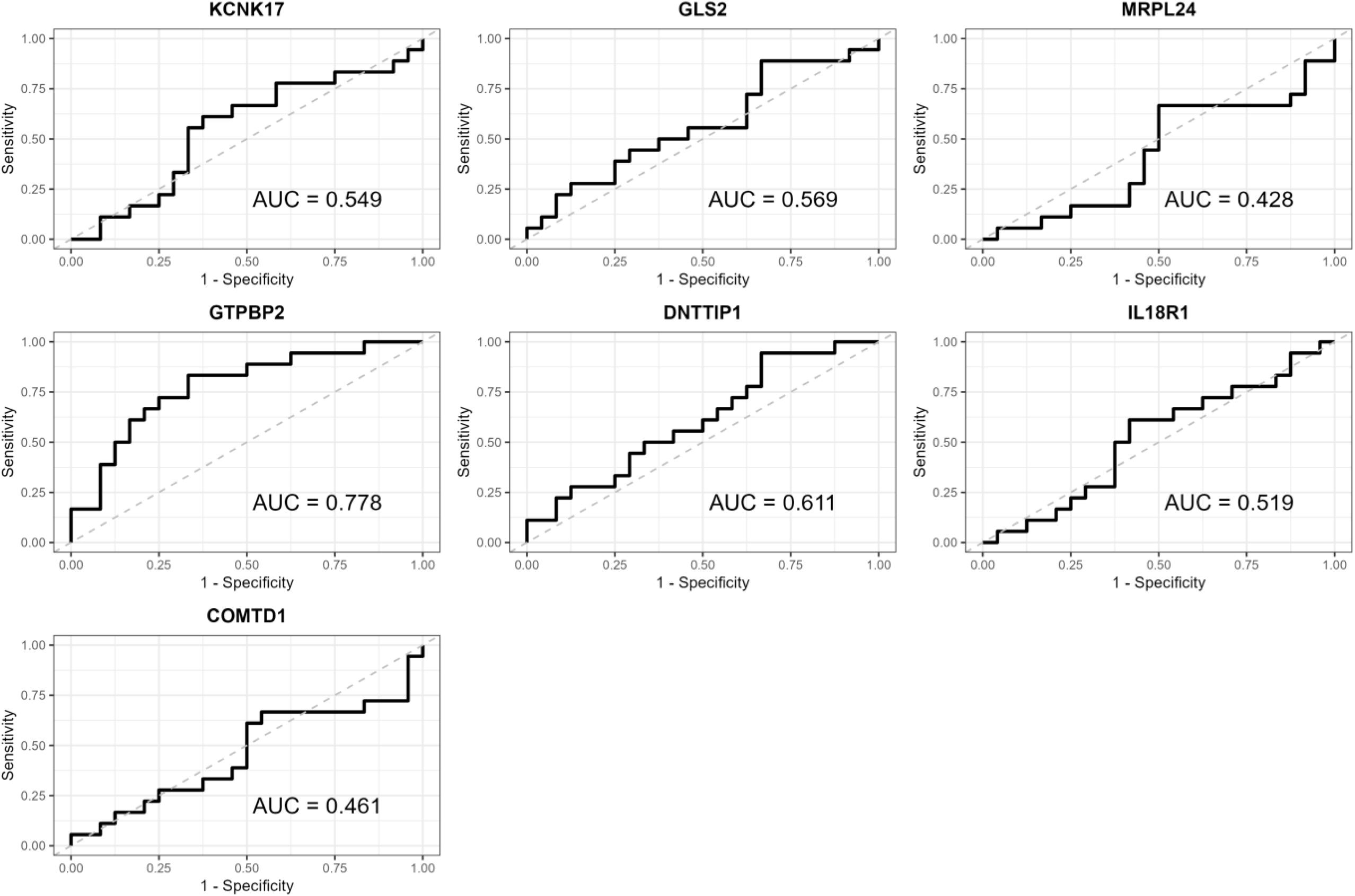
Receiver operating characteristic curves of each gene in the GSE33377 dataset.

## REFERENCES

1. Firestein GS. Evolving concepts of rheumatoid arthritis. Nature. 2003;423(6937):356–61.

2. Sokka T, Kautiainen H, Möttönen T, Hannonen P. Work disability in rheumatoid arthritis 10 years after the diagnosis. J Rheumatol. 1999;26(8):1681–5.

3. Aletaha D. Precision medicine and management of rheumatoid arthritis. J Autoimmun. 2020;110:102405.

4. Lin YJ, Anzaghe M, Schülke S. Update on the Pathomechanism, Diagnosis, and Treatment Options for Rheumatoid Arthritis. Cells. 2020;9(4):880.

5. van Schouwenburg PA, Rispens T, Wolbink GJ. Immunogenicity of anti-TNF biologic therapies for rheumatoid arthritis. Nat Rev Rheumatol. 2013;9(3):164–72.

6. Hetland ML, Christensen IJ, Tarp U, Dreyer L, Hansen A, Hansen IT, et al. Direct comparison of treatment responses, remission rates, and drug adherence in patients with rheumatoid arthritis treated with adalimumab, etanercept, or infliximab: results from eight years of surveillance of clinical practice in the nationwide Danish DANBIO registry. Arthritis Rheum. 2010;62(1):22–32.

7. Castro-Santos P, Laborde CM, Díaz-Peña R. Genomics, proteomics and metabolomics: their emerging roles in the discovery and validation of rheumatoid arthritis biomarkers. Clin Exp Rheumatol. 2015;33(2):279–86.

8. van der Pouw Kraan TCTM, van Gaalen FA, Huizinga TWJ, Pieterman E, Breedveld FC, Verweij CL. Discovery of distinctive gene expression profiles in rheumatoid synovium using cDNA microarray technology: evidence for the existence of multiple pathways of tissue destruction and repair. Genes Immun. 2003;4(3):187–96.

9. Lindberg J, af Klint E, Catrina AI, Nilsson P, Klareskog L, Ulfgren AK, et al. Effect of infliximab on mRNA expression profiles in synovial tissue of rheumatoid arthritis patients. Arthritis Res Ther. 2006;8(6):R179.

10. van der Pouw Kraan TC, Wijbrandts CA, van Baarsen LG, Rustenburg F, Baggen JM, Verweij CL, et al. Responsiveness to anti-tumour necrosis factor alpha therapy is related to pre-treatment tissue inflammation levels in rheumatoid arthritis patients. Ann Rheum Dis. 2008;67(4):563–6.

11. Badot V, Galant C, Nzeusseu Toukap A, Theate I, Maudoux AL, Van den Eynde BJ, et al. Gene expression profiling in the synovium identifies a predictive signature of absence of response to adalimumab therapy in rheumatoid arthritis. Arthritis Res Ther. 2009;11(2):R57.

12. Lindberg J, Wijbrandts CA, van Baarsen LG, Nader G, Klareskog L, Catrina A, et al. The gene expression profile in the synovium as a predictor of the clinical response to infliximab treatment in rheumatoid arthritis. PLoS One. 2010;5(6):e11310.

13. Toonen EJM, Gilissen C, Franke B, Kievit W, Eijsbouts AM, den Broeder AA, et al. Validation study of existing gene expression signatures for anti-TNF treatment in patients with rheumatoid arthritis. PLoS One. 2012;7(3):e33199.

14. Dennis G, Holweg CTJ, Kummerfeld SK, Choy DF, Setiadi AF, Hackney JA, et al. Synovial phenotypes in rheumatoid arthritis correlate with response to biologic therapeutics. Arthritis Res Ther. 2014;16(2):R90.

15. Aterido A, Cañete JD, Tornero J, Blanco F, Fernández-Gutierrez B, Pérez C, et al. A Combined Transcriptomic and Genomic Analysis Identifies a Gene Signature Associated With the Response to Anti-TNF Therapy in Rheumatoid Arthritis. Front Immunol. 2019;10:1459.

16. Oliver J, Nair N, Orozco G, Smith S, Hyrich KL, Morgan A, et al. Transcriptome-wide study of TNF-inhibitor therapy in rheumatoid arthritis reveals early signature of successful treatment. Arthritis Res Ther. 10 de marzo de 2021;23(1):80.

17. Tao W, Concepcion AN, Vianen M, Marijnissen ACA, Lafeber FPGJ, Radstake TRDJ, et al. Multiomics and Machine Learning Accurately Predict Clinical Response to Adalimumab and Etanercept Therapy in Patients With Rheumatoid Arthritis. Arthritis Rheumatol. 2021;73(2):212–22.

18. Hedman ÅK, Winter E, Yoosuf N, Benita Y, Berg L, Brynedal B, et al. Peripheral blood cellular dynamics of rheumatoid arthritis treatment informs about efficacy of response to disease modifying drugs. Sci Rep. 2023;13(1):10058.

19. Love MI, Huber W, Anders S. Moderated estimation of fold change and dispersion for RNA-seq data with DESeq2. Genome Biol. 2014;15(12):550.

20. Yu G, Wang LG, Han Y, He QY. clusterProfiler: an R package for comparing biological themes among gene clusters. OMICS. 2012;16(5):284–7.

21. Fernández Martínez JL, Fernández Muñiz MZ, Tompkins MJ. On the topography of the cost functional in linear and nonlinear inverse problems. GEOPHYSICS. 2012;77(1):W1–15.

22. Fernández-Martínez JL, Fernández-Muñiz Z, Pallero JLG, Pedruelo-González LM. From Bayes to Tarantola: New insights to understand uncertainty in inverse problems. Journal of Applied Geophysics. 2013;98:62–72.

23. deAndrés-Galiana EJ, Fernández-Martínez JL, Sonis ST. Sensitivity analysis of gene ranking methods in phenotype prediction. J Biomed Inform. 2016;64:255–64.

24. Deandrés-Galiana EJ, Fernández-Martínez JL, Saligan LN, Sonis ST. Impact of Microarray Preprocessing Techniques in Unraveling Biological Pathways. J Comput Biol. 2016;23(12):957–68.

25. Cernea A, Fernández-Martínez JL, deAndrés-Galiana EJ, Fernández-Ovies FJ, Fernández-Muñiz Z, Alvarez-Machancoses O, et al. Sampling Defective Pathways in Phenotype Prediction Problems via the Fisher’s Ratio Sampler. En: Rojas I, Ortuño F, editores. Bioinformatics and Biomedical Engineering. Cham: Springer International Publishing; 2018. p. 15–23.

26. Wickham H, Averick M, Bryan J, Chang W, McGowan LDa, François R, et al. Welcome to the Tidyverse. Journal of Open Source Software. 2019;4(43):1686.

27. Robin X, Turck N, Hainard A, Tiberti N, Lisacek F, Sanchez JC, et al. pROC: an open-source package for R and S+ to analyze and compare ROC curves. BMC Bioinformatics. 2011;12:77.

28. Wickham H. ggplot2: Elegant Graphics for Data Analysis. New York, NY: Springer; 2009.

29. Wickham H, François R, Henry L, Müller K, Vaughan D, Software P, et al. dplyr: A Grammar of Data Manipulation. 2023.

30. Kuhn M. Building Predictive Models in R Using the caret Package. Journal of Statistical Software. 2008;28(5):1–26.

31. Miller TL based on F code by A. leaps: Regression Subset Selection. 2024.

32. Pedersen TL. patchwork: The Composer of Plots. 2024.

33. Kassambara A. ggpubr: "ggplot2" Based Publication Ready Plots. 2023.

34. Fang J, Cao T, Liu C, Wang D, Zhang H, Tong J, et al. Association between magnesium, copper, and potassium intakes with risk of rheumatoid arthritis: a cross-sectional study from National Health and Nutrition Examination Survey (NHANES). BMC Public Health. 2023;23(1):2085.

35. Kang R, Kroemer G, Tang D. The tumor suppressor protein p53 and the ferroptosis network. Free Radic Biol Med. 2019;133:162–8.

36. Zhao T, Yang Q, Xi Y, Xie Z, Shen J, Li Z, et al. Ferroptosis in Rheumatoid Arthritis: A Potential Therapeutic Strategy. Front Immunol. 2022;13:779585.

37. Hisada R, Yoshida N, Orite SYK, Umeda M, Burbano C, Scherlinger M, et al. Role of Glutaminase 2 in Promoting CD4+ T Cell Production of Interleukin-2 by Supporting Antioxidant Defense in Systemic Lupus Erythematosus. Arthritis Rheumatol. 2022;74(7):1204–10.

38. Chu JY, McCormick B, Vermeren S. Small GTPase-dependent regulation of leukocyte-endothelial interactions in inflammation. Biochem Soc Trans. 19 de junio de 2018;46(3):649–58.

39. Wang X, Li W, Lou N, Han W, Hai B, Xiao W, et al. High Expression of DNTTIP1 Predicts Poor Prognosis in Clear Cell Renal Cell Carcinoma. Pharmgenomics Pers Med. 2023;16:1–14.

40. Landy E, Carol H, Ring A, Canna S. Biological and clinical roles of IL-18 in inflammatory diseases. Nat Rev Rheumatol. 2024;20(1):33–47.

41. Li J, Liu J, Li J, Feng A, Nie Y, Yang Z, et al. A risk prognostic model for patients with esophageal squamous cell carcinoma basing on cuproptosis and ferroptosis. J Cancer Res Clin Oncol. 2023;149(13):11647–59.

42. Di Nottia M, Marchese M, Verrigni D, Mutti CD, Torraco A, Oliva R, et al. A homozygous MRPL24 mutation causes a complex movement disorder and affects the mitoribosome assembly. Neurobiol Dis. 2020;141:104880.

43. Gopisetty G, Thangarajan R. Mammalian mitochondrial ribosomal small subunit (MRPS) genes: A putative role in human disease. Gene. 2016;589(1):27–35.

44. Lin S, Hannon E, Reppell M, Waring JF, Smaoui N, Pivorunas V, et al. Whole Blood DNA Methylation Changes Are Associated with Anti-TNF Drug Concentration in Patients with Crohn’s Disease. J Crohns Colitis. 2024;18(8):1190–201.

45. Ek WE, Karlsson T, Höglund J, Rask-Andersen M, Johansson Å. Causal effects of inflammatory protein biomarkers on inflammatory diseases. Sci Adv. 2021;7(50):eabl4359.

46. Eriksson C, Engstrand S, Sundqvist KG, Rantapää-Dahlqvist S. Autoantibody formation in patients with rheumatoid arthritis treated with anti-TNF alpha. Ann Rheum Dis. 2005;64(3):403–7.

47. Ibáñez-Costa A, Perez-Sanchez C, Patiño-Trives AM, Luque-Tevar M, Font P, Arias de la Rosa I, et al. Splicing machinery is impaired in rheumatoid arthritis, associated with disease activity and modulated by anti-TNF therapy. Ann Rheum Dis. 2022;81(1):56–67.

